# Combining Two Classes of Epigenetic Modifiers and Their Effect on Prostate Cancer Cells

**DOI:** 10.1101/2020.12.01.404574

**Authors:** Mudassir K. Lodi, Ekaterine Goliadze, Masoud H. Manjili, Georgi Guruli

**Affiliations:** Virginia Commonwealth University, Nothing to disclose; Department of Surgery, Division of Urology, Virginia Commonwealth University School of Medicine, Nothing to disclose; Department of Microbiology and Immunology, Virginia Commonwealth University School of Medicine; Department of Surgery, Division of Urology, Virginia Commonwealth University School of Medicine

**Author notes:** Corresponding Author: Georgi Guruli, MD, PhD, Nothing to disclose.

## Abstract

The main objective of this experiment was to determine and study the effects of combining two epigenetic modifiers, 5-azacyticidine (5-AzaC) and SB939, on a RM-1 murine prostate cancer cell model. The effectiveness of this combination on prostate cancer cells has not been previously studied. The study was implemented on ex vivo cell models to gain a better understanding of the true effects of the combination therapy on prostate cancer cells. Two variations of the combination therapy were tested in this study, each with different concentrations of SB939 (100nm and 200nm).

To determine the effectivity of the combination therapy on prostate cancer cells, three factors were measured: cell proliferation, cancer testis antigen (CTA) expression, and apoptosis rate. To measure cell proliferation, a cell proliferation assay was conducted, and absorption rate was measured through a 450 nm wavelength. CTA expression was measured through a quantitative polymerase chain reaction (quant-PCR). For this study, the expression rates of five CTAs were measured (TEX15, CEP55, CCNA1, P1A, SPA17). Apoptosis rate was measured through an Annexin-V assay, in which two markers, Annexin-V and 7-AAD, were used.

We found that SB939 combined with 5-AzaC show highest efficacy compare to each drug alone in terms of inhibiting tumor cell proliferation, as well as inducing tumor cells apoptosis and enhancing tumor cell immunogenicity by the induction of the expression of CTAs. This combination proved to be effective in combating murine prostate cancer cells, and can potentially be effective within in vivo models due to its high toxicity to these cancer cells, and its ability to render prostate cancer more immunogenic.

## Introduction

Despite advancements in the medicine and cancer treatment in particular, cancer is the second leading cause of death in the United States (https://www.medicalnewstoday.com/articles/282929). Prostate cancer is the most prevalent form of cancer in men, with an estimated 191, 930 new cases and expected 33, 330 deaths in 2020 ^1^. Androgen deprivation therapy is the main treatment option for advanced prostate cancer cases. However, it is not curative, and once it fails, treatment options become limited. The need for the development of the new therapeutic modalities is imperative, and several treatment options have been tested recently. Immunotherapy is one of the most common explored treatment modalities ^2, 3^, but its effectiveness in the treatment of the advanced prostate cancer seems to be limited. Both the low immunogenicity of prostate cancer and possible suppression of the immune system play a significant role in the decreased effectiveness of the immunotherapy. To overcome this problem, we explore the value of epigenetic modification, which should increase the immunogenicity of prostate cancer and immunomodulation.

Many drug therapies specifically target tumors, the masses of proliferated cells that grow during various stages of cancer. One emerging class of therapies involve epigenetic modifiers, genes that directly modify the epigenome through one of DNA methylation, post-translational modification of chromatin, and alteration to chromatin structure. There is evidence to suggest that epigenetic modifiers have a profound effect on the growth and proliferation of tumors. Cancer therapies have also increasingly utilized immunomodulation, defined as the modification of the immune system. These immunotherapies typically utilize immunomodulators to effect these changes.

One class of epigenetic modifiers, DNA methyltransferase (DNMT) inhibitors, are currently used to treat several forms of cancer, including myelodysplastic syndrome (MDS) and acute myeloid leukemia (AML). They are one of the more prominent epigenetic modifier treatments commercially available. One such DNMT, 5-azacytidine, when combined with immunomodulation, has been shown to decrease the production of cancer-testis antigens (CTA) within a murine prostate cancer model (*ex vivo*), and decrease malignant tumor cell growth ^4^.

Histone Deacetylase Inhibitors (HDIs), a different type of epigenetic modifier, are a class of cytostatic agents that inhibit the proliferation of tumor cells in culture and in vivo by inducing cell cycle arrest, differentiation and / or apoptosis. HDIs have been clinically proven to be effective in treating cancers of the lymphatic system, such as cutaneous T-cell lymphoma, as well as parasitic and inflammatory diseases ^5^. However, no such studies have been done to suggest it as practical treatment method in prostate cancer. Although HDIs have been evaluated as a treatment option itself, they have never been coupled with other epigenetic modifiers and used as a treatment drug for any type of cancer or disease.

SB939 (pracinostat), a HDI, is currently used to treat cancers such as T-cell lymphoma and acute myelocytic leukemia (AML). SB939 has been clinically proven to increase survival rates in elderly AML patients ^6^. In another study, SB939 was shown to have anticancer properties, and was shown to have maximum efficacy in doublet therapy. In addition, pracinostat shows these same results against more therapeutically challenging cancers ^7^. SB939 was also cited as a viable treatment drug for acute myeloid leukemia based on the studies discussed above ^8^.

With innovations in precision medicine, there is an active interest in the benefits of combination therapies. It will beneficial to study the effects on prostate cancer cells with a combination of epigenetic modifiers. The HDI drug SB939 will be tested and combined with the DNMT drug 5-azacytidine. Because of efficiency of both SB939 and 5-azacytidine in treating acute myeloid leukemia, and the specific effects of 5-azacytidine on CTAs within prostate cancer cells, it can be reasonably hypothesized that a combination of the two drugs would reduce the tumorgenicity of prostate cancer cells in an ex vivo setting.

Because of its prominence in men in the United States and around the world, it would be beneficial to study some alternative treatment methods for prostate cancer.

Studying the effects of this combination therapy on prostate cancer specifically can open a wide range of possibilities. If it is found that the combination of pracinostat with 5-azacytidine results in a lower tumor immunogenicity than existing treatment drugs in the market, other combinations can also be tested for their effects, and the most successful treatment could be implemented into the market. Although there are several treatment drugs in the market for prostate cancer already, it would be beneficial to continually keep discovering new and more efficient therapies. Studying combinations between various classes of epigenetic modifiers could potentially broaden the scope of treatment drugs.

## Methods

### Prostate Cancer Cell Line

RM-1 cell line is an androgen-independent murine prostate cancer cell line. It was a gift from Dr. Timothy C. Thompson (Baylor College of Medicine, Houston, TX). This model was generated by transduction of cells with the ras and myc oncogenes, yielding a poorly differentiated adenocarcinoma. Tumor cells are maintained in complete media at 37°C in 5% CO2. For in vivo studies, RM-1 cells (50,000 cells/100 μl) will be inoculated subcutaneously (SC) into the right shaved flank of C57BL/6 mice, and tumor establishment will be determined by palpation. Tumor growth will be assessed every other day by microcalipers.

### Study compounds

5-azacitidine and SB939 were obtained from Santa Cruz Biotechnology, Inc. (Dallas, Texas). 5-azacitidine was dissolved in 100% dimethylsulfoxide (DMSO) (Sigma-Aldrich, St. Louis, MO) before further dilution in cell culture media. SB939 was dissolved in 100% DMSO as well, at 10mM concentration. Final DMSO concentrations will be kept at a constant 0.1% for all samples, including controls, unless otherwise stated. Both drugs were maintained as stock solutions for *in vitro* experiments at −20°C for no longer than 1 month.

### Cell Proliferation Assay

Cell proliferation was measured using a colorimetric assay with WST-1 reagent, which quantifies mitochondrial metabolic activity of viable cells per manufacturer’s instructions (Roche, Indianapolis, IN). Cells were cultured in 96-well microplates in a concentration of 5×10^4^ cells/mL (in RPMI with 10% FBS) and cultivated for 48 h in a humidified atmosphere (37.0°C; 5% CO2). After 44 h, 10 μL of WST-1 were added and cells were incubated for an additional 4 h. During this incubation period, viable cells converted WST-1 to a water soluble formazan dye. Cell viability was measured at 450 nm in a microplate reader (Bio Rad) (Reference wavelength: 655 nm).

### Annexin V Assay

RM-1 cells were collected and double stained with FITC-conjugated annexin V and 7-AAD, according to the manufacturer’s instructions (Biolegend, San Diego, CA). Briefly, cells were collected after 48 hours of culture under different conditions, washed twice with cold Biolegend’s Cell Staining buffer, and resuspended in Annexin V Binding Buffer at a concentration of 10^5^ cells/ml. Cells were incubated with Annexin V-FITC and 7-AAD for 15 minutes at room temperature. After washing steps all samples were analyzed within 30 min. Data were acquired using a BD FACSCANTO II benchtop analyzer (Becton Dickinson, San Jose, CA) and analysis was performed using BD FACSDiva software (BD) and FCS Express (De novo software, Los Angeles, CA).

### RNA Isolation

Cells were grown in 6-well plates, and collected once they were almost confluent, and used immediately for RNA isolation. RNA was routinely isolated by TRIzol (Invitrogen, Carlsbad, CA). For that, tissue was placed in cold TRIzol (4°C) and immediately homogenized using a Bio-Gen PRO200 Homogenizer (Pro scientific Inc., Oxford, CT) The protocol for TRIzol isolation was then completed following the manufacturer’s instructions. Isolated RNA was purified with an RNeasy Mini Kit (Qiagen, Valencia, CA). RNA yields were determined spectrophotometrically at 260 nm.

### Quantitative Real Time Polymerase Chain Reaction (qRT-PCR)

After RNA isolation, cDNA was synthesized using the ThermoScript RT-PCR System (Invitrogen) from 1mg of total RNA using Random Primer. In each experiment, at least three independent reactions were performed to obtain the mean. QRT-PCR was performed in triplicate including a non-template control using the Mx3000P system (Agilent Technologies, Inc., Santa Clara, CA). Oligos (Invitrogen) were designed using Primer3 software (White head Institute of Biomedical Research MIT. Boston, MA), and are presented in Table I. GAPDH and P1A Primers were obtained from Qiagen.

**Table I.**
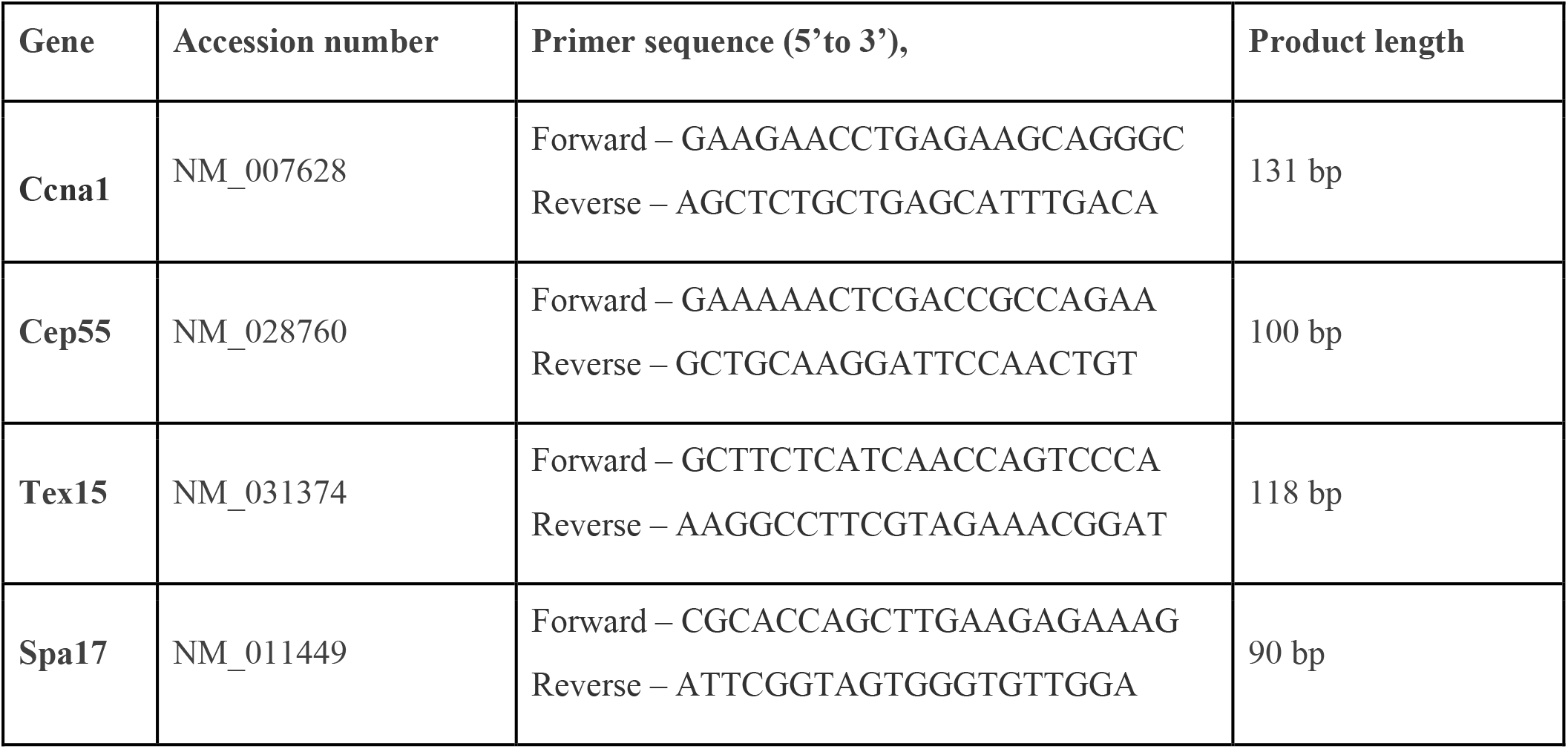
Primers for polymerase chain reaction (PCR)

Real-time RT-PCR reactions were performed in 20ml volumes with 10 ml of SensiFAST SYBR Lo-ROX Kit (Bioline, Taunton, MA) 2ml of cDNA template and 0.5ml each of the forward and reverse primers of the gene of interest (GOI). The cDNA used for the PCR reactions was diluted 1:35 for each GOI. PCR conditions were as follows: an initial denaturation step (10 min at 95°C), 40 cycles consisting of three steps (30 sec at 95°C, 1 min at 55°C, 30 sec at 72°C), and 1 cycle consisting of three steps (1 min at 5°C, 30 sec at 55°C, and 30 sec at 95°C). The cycle threshold (CT) value was the PCR cycle number in which the fluorescence signal was significantly distinguishable from the baseline for the first time.

The housekeeping gene (GAPDH) was used as an endogenous control for target gene expression evaluation. Expression values of each gene were normalized to the expression of GAPDH of a given sample. Data were presented by the relative amount of mRNA with the formula 2 DDCT, which stands for the difference between the CT of a gene of interest and the CT of the housekeeping gene (GAPDH).

#### Study design – ex vivo

RM-1 murine prostate cancer cells were exposed to 5-azacytidine (0.5 μM concentration) and / or SB939 (100nm and 200nm concentrations). A DMSO control and one group with no treatment will be included as well. After 24 and 48 hours of exposure, cells were collected, and cell proliferation assay and Annexin V apoptosis assay were performed. For western blot, cells were collected after their exposure to 5-azacytidine and SB939 as it was described before, protein was extracted and the expression of the acH3 was evaluated.

### Statistical analysis

For a single comparison of 2 groups, the Student *t* test was used (SigmaPlot Software; SPSS, Chicago, IL). If data distribution was not normal, the Mann-Whitney rank sum test was run instead. A z-test was performed to evaluate the significance of differences between the experimental groups in the flow cytometry assays when discrete data were presented. For all analysis, the level of significance was set at a probability of 0.05 to be considered significant. Data are presented as the mean ± standard error of the mean (SEM).

## Results

### SB939 combined with 5-AzaC inhibits tumor cell proliferation with the highest efficiency

RM-1 cells in suspension were exposed to: (i) 5-AzaC (1μM concentration); (ii) SB939 (100 nM); (iii) SB939 (200nM); (iv) 5-AzaC + SB939 (100nM); (v) 5-AzaC + SB939 (200nM). Cells treated with DMSO provided the control. Results are presented in table 2 and figure 1. While the use of 5-AzaC did not affect cell proliferation significantly, it seems that any use of SB939 depressed RM-1 cells proliferation significantly (P=0.031 for SB939, 100nM concentration, with CI of 0.0660 to 1.244; P=0.017 and CI was 0.147 to 1.252 for SB939 200nM solution; P=0.001 and CI – 0.449 to 1.507 for the combination of 5-AzaC and SB939, 100nM; and P=0.004 with CI of 0.342 to 1.432 for the combination of 5-AzaC and SB939, 200nM) in comparison to control (DMSO). There was no difference between different concentrations of SB939.

**Table 2.** Cell Proliferation Assay (Absorbance Rate)

| **Drug Therapy** | **Absorbance** |
| --- | --- |
| **DMSO** | **1.286 ± 0.25** |
| **5-AzaC 1μM** | **0.984 ± 0.34** |
| **SB 100 nm** | **0.631 ± 0.12** |
| **SB 200 nm** | **0.586 ± 0.08** |
| **5-AzaC + 100 nm** | **0.308 ± 0.03** |
| **5-AzaC + 200 nm** | **0.399 ± 0.07** |

**Figure 1.**
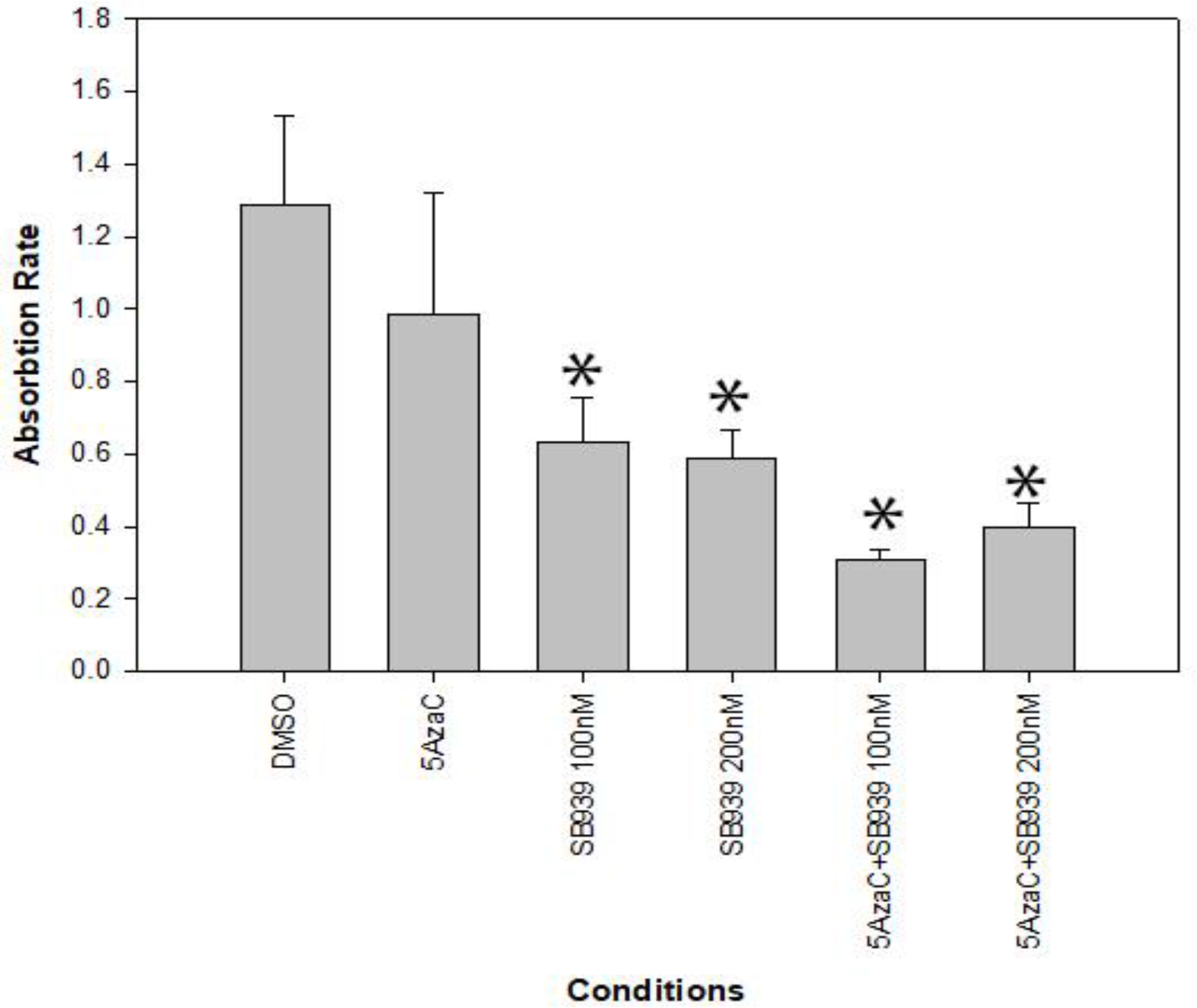
RM-1 cell proliferation assay. RM-1 were exposed to: (i) 5-AzaC (1μM concentration); (ii) SB939 (100 nM); (iii) SB939 (200nM); (iv) 5-AzaC + SB939 (100nM); (v) 5-AzaC + SB939 (200nM). Cells treated with DMSO provided the control. Cell proliferation was measured using a colorimetric assay with WST-1 reagent, which quantifies mitochondrial metabolic activity of viable cells per manufacturer’s instructions (Roche, Indianapolis, IN). Cells were cultured in 96-well microplates in a concentration of 5×104 cells/mL (in RPMI with 10% FBS) and cultivated for 48 h in a humidified atmosphere (37.0°C; 5% CO2). Cell viability was measured at 450 nm in a microplate reader (Bio Rad) (Reference wavelength: 655 nm). The * represents statistically significant results from the DMSO.

### SB939 promotes 5-AzaC-induced apoptosis in tumor cells

We utilized Annexin V / 7-ADD staining to establish apoptotic capabilities of the 5-AzaC and SB939 concerning the RM-1 cells. There were 5 treatment groups for RM-1 cells: (i) 5-AzaC (1μM concentration); (ii) SB939 (100 nM); (iii) SB939 (200nM); (iv) 5-AzaC + SB939 (100nM); 5-AzaC + SB939 (200nM). Cells treated with DMSO provided the control. Results are presented in Table 3 and Figures 2 and 3. Apoptosis was increased in comparison to control by using 5-AzaC alone (P<0.001, CI: −19.094 to −11.526). SB939 did not increase RM-1 apoptosis alone, and addition of the SB939 to 5-AzaC at a concentration of 100nM did not cause further increase in apoptosis. However, addition of 200nM concentration of SB939 to the 5-AzaC resulted in the further increase of the apoptosis in comparison to 5-AzaC alone (p=0.007, CI: - 28.922 to −8.492).

**Table 3.**
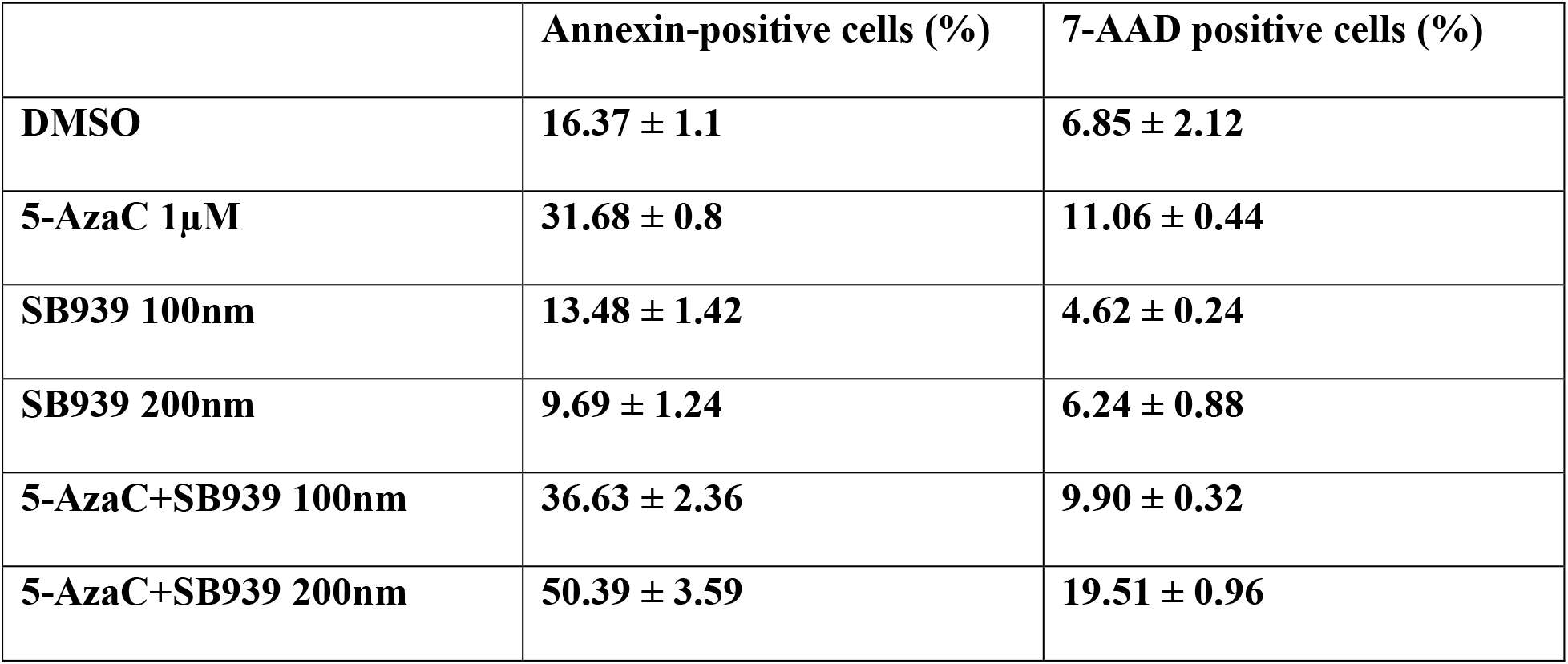
RM-1 cells apoptosis rate.

|  | **Annexin-positive cells (%)** | **7-AAD positive cells (%)** |
| --- | --- | --- |
| **DMSO** | **16.37 ± 1.1** | **6.85 ± 2.12** |
| **5-AzaC 1μM** | **31.68 ± 0.8** | **11.06 ± 0.44** |
| **SB939 100nm** | **13.48 ± 1.42** | **4.62 ± 0.24** |
| **SB939 200nm** | **9.69 ± 1.24** | **6.24 ± 0.88** |
| **5-AzaC+SB939 100nm** | **36.63 ± 2.36** | **9.90 ± 0.32** |
| **5-AzaC+SB939 200nm** | **50.39 ± 3.59** | **19.51 ± 0.96** |

**Figure 2.**
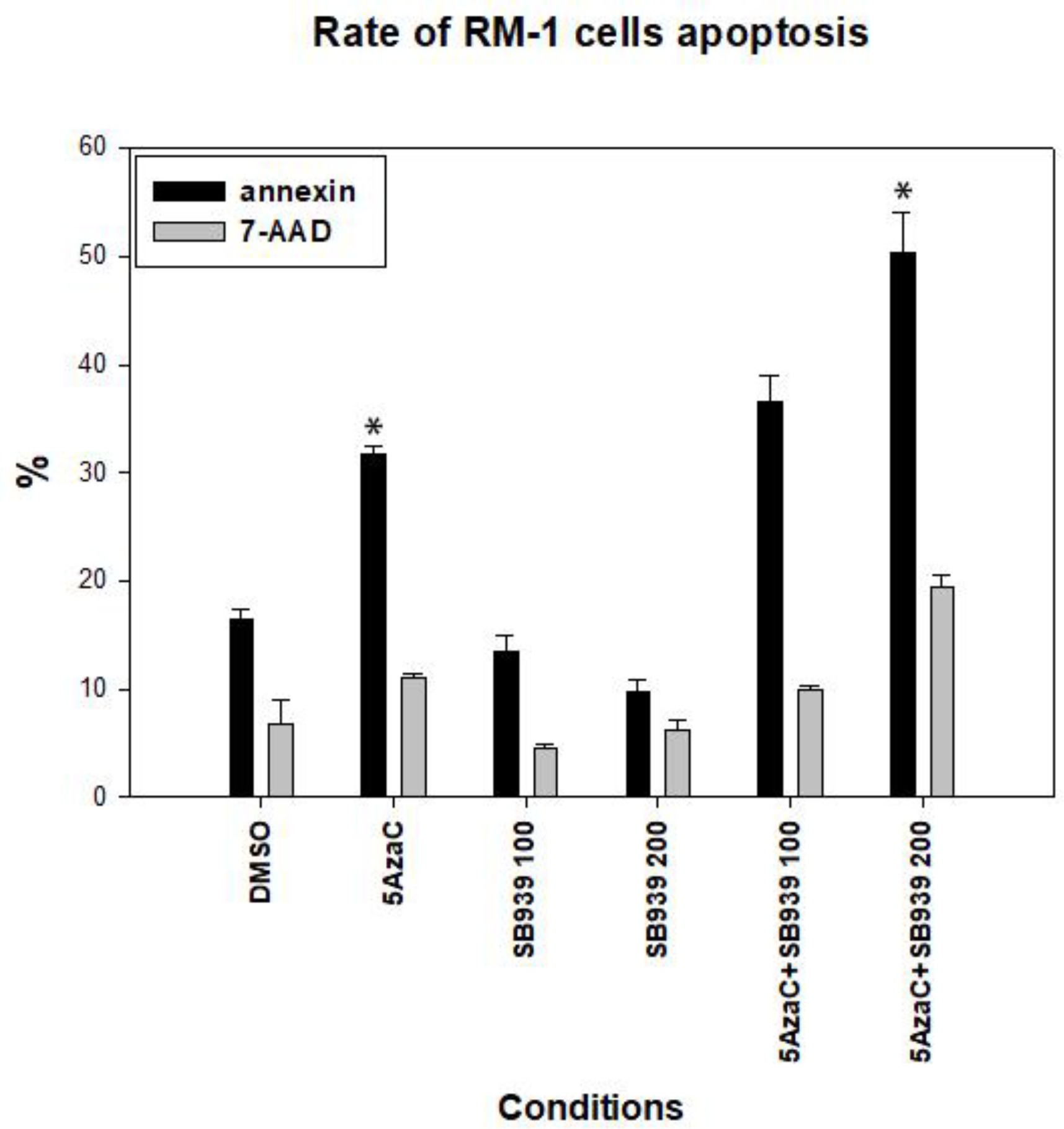
Rate of RM-1 cells apoptosis after exposing them to the 5-AzaC, SB939 and there combinations. Apoptosis rate was assessed by Annexin V / 7-ADD staining of the RM-1 cells. There were 5 treatment groups for RM-1 cells: (i) 5-AzaC (1μM concentration); (ii) SB939 (100 nM); (iii) SB939 (200nM); (iv) 5-AzaC + SB939 (100nM); (v) 5-AzaC + SB939 (200nM). Cells were incubated with Annexin V-FITC and 7-AAD for 15 minutes at room temperature. After washing steps all samples were analyzed within 30 min. Data were acquired using a BD FACSCANTO II benchtop analyzer (Becton Dickinson, San Jose, CA) and analysis was performed using BD FACSDiva software (BD) and FCS Express (De novo software, Los Angeles, CA). The * represents statistically significant results from the DMSO.

**Figure 3.**
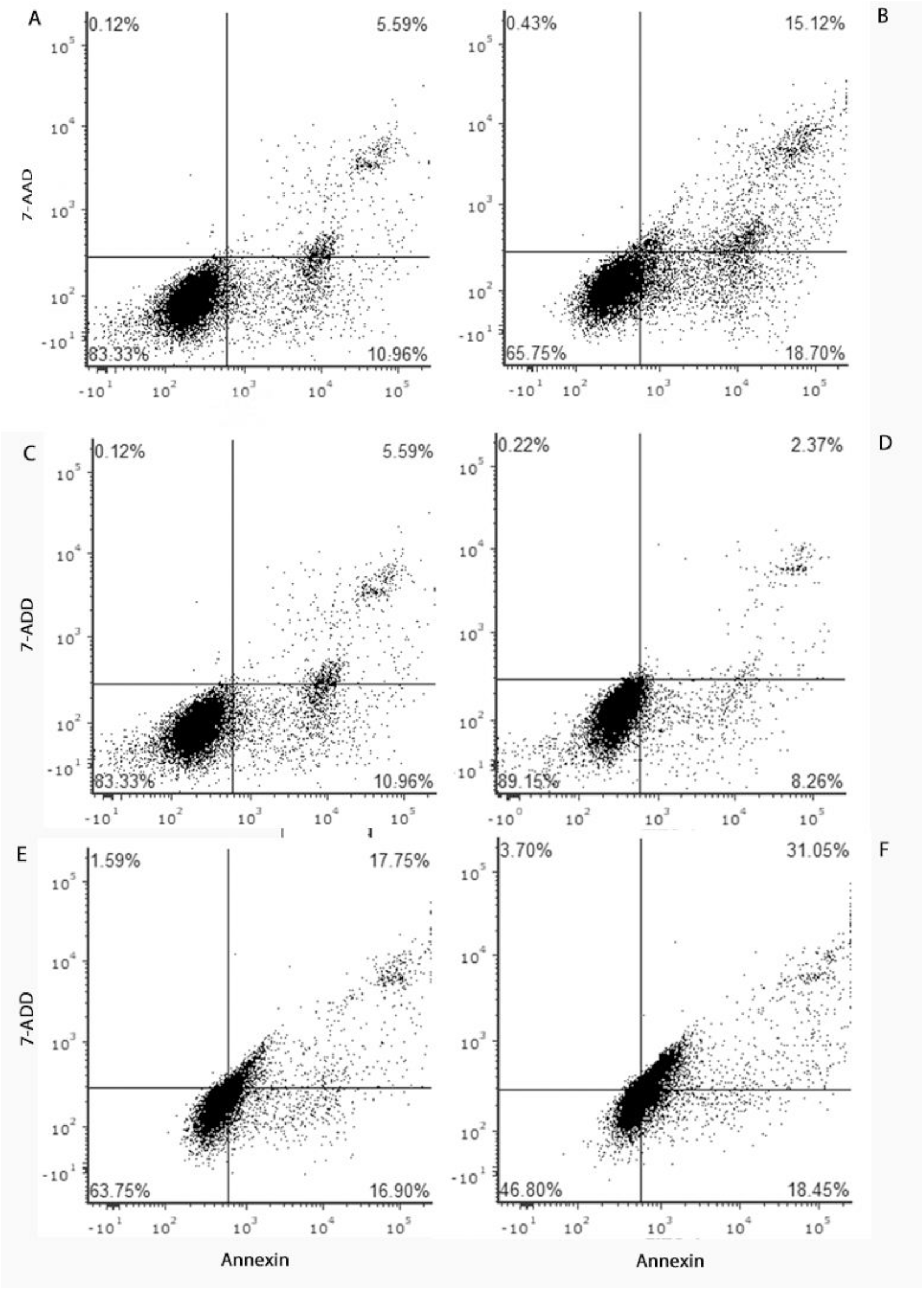
Rate of RM-1 cells apoptosis after exposing them to the 5-AzaC, SB939 and their combinations, by Annexin V assay. Representative flow cytometry dot plot images are shown. A: DMSO (control); B: 5-AzaC (1μM concentration); C: SB939 (100 nM); D: SB939 (200nM); E: 5-AzaC + SB939 (100nM); F: 5-AzaC + SB939 (200nM). Data were acquired using a BD FACSCANTO II benchtop analyzer (Becton Dickinson, San Jose, CA) and analysis was performed using BD FACSDiva software (BD) and FCS Express (De novo software, Los Angeles, CA).

### SB939 enhances the 5-AzaC-induced expression of CTA in tumor cells

We already established the ability of 5-AzaC to increase the expression of certain CTAs in the prostate cancer cells (RM-1 cells)(Citation – sulek). For this work, we picked 5 CTAs and evaluated the possible influence of the SB939 on their expression, alone or in combination with 5-AzaC. There were 5 experimental groups, as in previous experiments: (i) 5-AzaC (1μM concentration); (ii) SB939 (100 nM); (iii) SB939 (200nM); (iv) 5-AzaC + SB939 (100nM); (v) 5-AzaC + SB939 (200nM). Results are presented in Table 4 and Figure 4. As it was expected, 5- AzaC induced significant elevation of the expression of all 5 CTAs. SB939 alone had no influence on the expression of the CTAs. However, when combined with 5-AzaC, at the SB939 concentration of 100nM, there was a further increase in the expression of all 5 CTAs tested, and that increase was statistically significant. Interestingly, addition of SB939 at a concentration 200nM had no additive effect.

**Table 4.** Expression of different CTAs by murine prostate cancer cells.

|  | **CCNA1** | **CEP55** | **TEX15** | **P1A** | **SPA17** |
| --- | --- | --- | --- | --- | --- |
| **5AzaC** | **3.74 ± 0.143** | **4.71 ± 0.187** | **1.87 ± 0.088** | **3.047 ± 0.034** | **2.743 ± 0.0464** |
| **SB939 100** | **1.263 ± 0.074** | **1.52 ± 0.076** | **1.158 ± 0.082** | **1.083 ± 0.0165** | **1.627 ± 0.094** |
| **SB939 200** | **0.083 ± 0.012** | **0.41 ± 0.0721** | **0.59 ± 0.15** | **0.0277 ± 0.0118** | **0.717 ± 0.0754** |
| **5AzaC+SB939 100nm** | **5.213 ± 0.122** | **5.93 ± 0.0436** | **3.47 ± 0.07** | **4.4 ± 0.171** | **3.53 ± 0.112** |
| **5AzaC+SB939 200nm** | **2.067 ± 0.0524** | **3.223 ± 0.103** | **0.89 ± 0.186** | **1.707 ± 0.0865** | **1.413 ± 0.096** |

**Figure 4.**
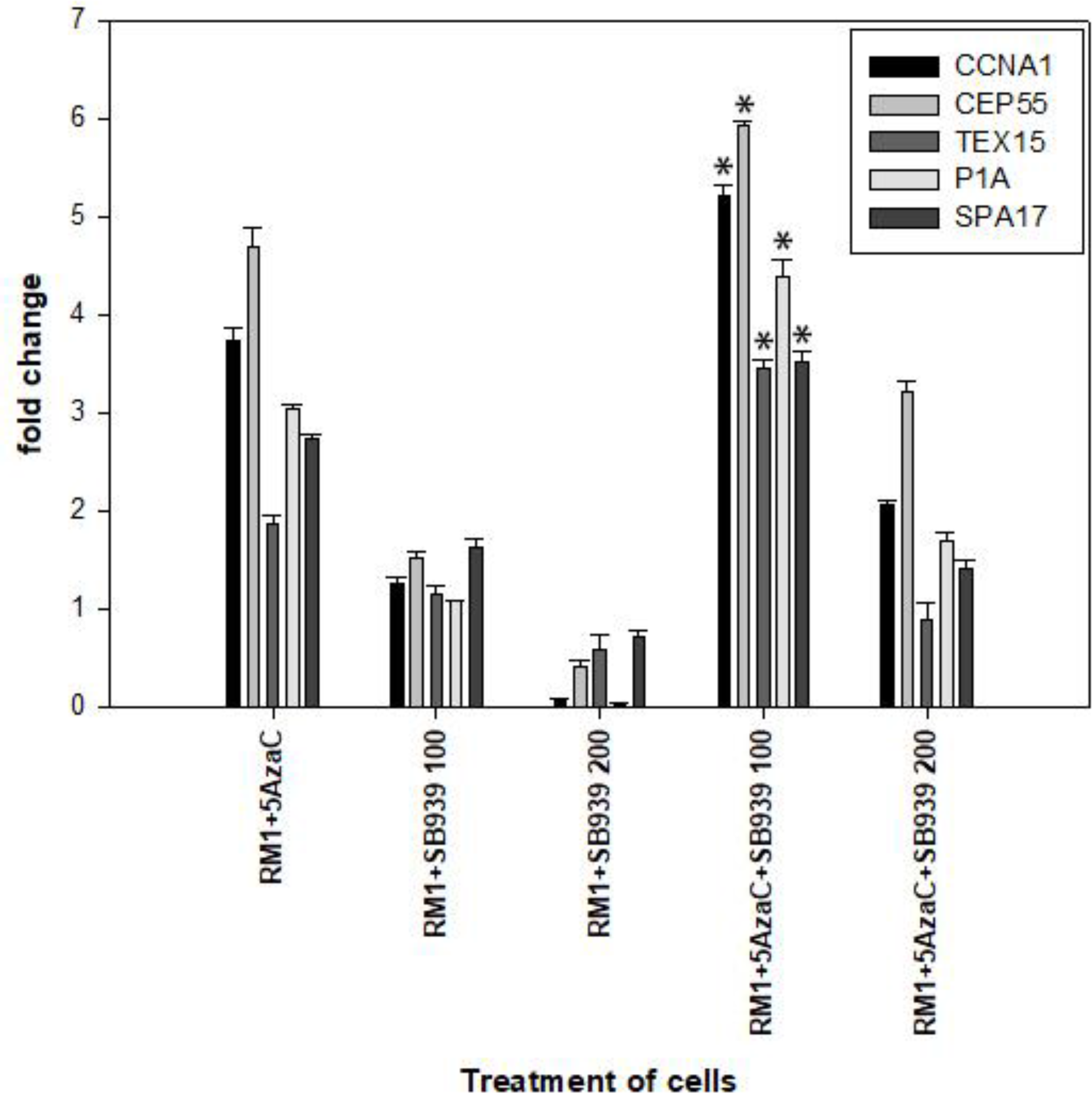
Expression of various CTAs by murine prostate cancer cells (RM-1 cells). There were 5 experimental groups, as in previous experiments: (i) 5-AzaC (1μM concentration); (ii) SB939 (100 nM); (iii) SB939 (200nM); (iv) 5-AzaC + SB939 (100nM); (v) 5-AzaC + SB939 (200nM). The housekeeping gene (GAPDH) was used as an endogenous control for target gene expression evaluation. Expression values of each gene were normalized to the expression of GAPDH of a given sample. Data were presented by the relative amount of mRNA with the formula 2 DDCT, which stands for the difference between the CT of a gene of interest and the CT of the housekeeping gene (GAPDH). The * represents statistically significant results from the group 1, treated with 5-AzaC only.

## Discussion

Upon analysis of the results, it can be concluded that the combination of the two epigenetic modifiers, SB939 and 5-AzaC, resulted in the increased apoptosis of the murine prostate cancer cells (RM-1 cells) in comparison to either modifier alone. the process of apoptosis, phosphatidylserine (PS) residues appear on the surface of the cell, and therefore can be used as an indicator of apoptosis rates. Because of their high affinity for phosphatidylserine, two biomarkers, annexin-V and 7-AAD, were chosen to stain the cells.

Upon completion of the annexin-V assay, data percentages of cells stained with either biomarker were collected and analyzed. The DMSO (control) resulted in 16.87% Annexin and 6.85% 7- AAD. Therefore, group percentage values above these two values would indicate success, in that the given therapy resulted in a higher apoptosis rate than normal conditions.

The components of the combination drug therapy were tested each as their own group. The 5-Azacytidine yielded 31.68% Annexin and 11.06% 7-AAD. SB939 100nm yielded 13.48% Annexin and 4.62% 7-AAD, while SB939 200nm yielded 9.69% Annexin and 6.24% 7-AAD. Both SB939 concentrations resulted in values lower than the DMSO, while 5-Azacytidine seemed to be more effective in inducing cell apoptosis.

The combination therapies, as hypothesized, did lead to higher percentages of cell apoptosis than the control and separate components. The 5-Azacytidine and SB939 100nm combination resulted in 36.63% Annexin and 9.90% 7-AAD. The 5-Azacytidine and SB939 200nm combination resulted in 50.39% Annexin and 19.51% 7-AAD, both values being significantly higher than all other group percentages.

Based on the results, it can be concluded that SB939 individually does not induce a higher apoptosis rate. 5-Azacytidine did induce higher Annexin and 7-AAD. However, the combination of these two components resulted in the highest Annexin and 7-AAD percentages of all the groups, leading to the conclusion that the combination therapy is effective in terms of inducing apoptosis.

According to a phase 2 study conducted in 2019, this combination therapy has been proven to be effective in older AML patients^9^. The study concludes that due to its efficacy in older patients, the combination of SB939 and 5-azacytidine could potentially be utilized as a viable AML treatment.

Another study also references the synergy between these two classes of epigenetic modifiers, and concluded that a combination of these two classes were found to induce gene re-expression, leading to an increased apoptosis rate^10^. This concurs with the results of the paper, in that it was also found that this combination therapy could have the same benefits in prostate cancer patients, based on the results of the ex vivo experiments.

Similar results indicating the effectiveness of the combination therapy in reducing tumor cell proliferation were found in the cell proliferation experiments as well. The cell proliferation experiments were measured by absorbance of a 450nm wavelength. In this conduction of the cell proliferation assay, lower absorbance indicate decreased cell viability. Lower cell viability rates compared to the control group show an increased effectiveness of the combination therapy.

The control group had an absorbance of 1.286, which, as expected, was the highest value of all the groups. The separate components each had a lower absorbance rate, with higher concentrations of SB939 resulting in lower absorbance rates. This is most likely due to the basic effects of the components themselves, in that administering more of the drug would naturally result in decreased proliferation in ex vivo models. The combination therapy had a significantly lower absorbance rate compared to the remainder of the groups, with the 5-azacytidine and SB939 100nm combination resulting in a slightly lower absorbance than the 5-azacytidine and SB939 200 nm combination.

Based on the results of the cell proliferation assay, it can be concluded that the combination therapy has an inverse effect on the proliferation of murine prostate cancer cells. This concurs with the data from the apoptosis rate, in that the combination therapy is toxic toward the viability of prostate cancer cells.

Cancer testis antigens play an important role in immunotherapy, as they act as targets for tumor-directed immune responses. Therefore, the level of expression of different cancer-testis antigens (CTA) in RM-1 cells was analyzed using quantitative PCR. As was hypothesized, 5-azacytidine induced increased expression of CTAs in the prostate cancer cells. Though SB939 alone did not affect CTAs expression significantly, the combination of SB939 (100nM) and 5-azacytidine had an additional increase in the expression of the given CTAs compared to 5-azacytidine alone (Table 4). That increase was not observed with the 200 nM concentration of the SB939. That can be because of the more toxic influence of the 200nM concentration of SB939, as we saw in apoptosis experiments. That might have limited the response to this combination.

Based on the results from the three assessments of the 5-azacytidine and SB939 combination therapy, it can be concluded that it is effective in inducing apoptosis and limiting cell proliferation, but would not be a viable immunotherapy solution. The results indicate that the therapy is effective in neutralizing prostate cancer cells.

